# Live confocal imaging of cellular touch responses upon local quantifiable mechanical stimulation in plant cells

**DOI:** 10.64898/2026.09.10.750548

**Authors:** A. Bellandi, C. Lionnet, D. Arico, N. German, M.O. Lenz, C. Kirchhelle, I. Fobis-Loisy, O. Hamant

## Abstract

Plant cells grow and differentiate in an ever-changing environment characterized by transient signals and stimuli. The ability of plant cells to perceive, integrate, and dynamically respond to these stimuli underpins a plant adaptation and survival.

Among the plethora of complex stimuli plant cells are exposed to, several stimuli have a mechanical component, for example, wind, touch, contact with insects, penetration of pathogens, and even intrinsic mechanical stresses arising during tissue growth.

Despite the presence of a load-bearing cell wall that separates cells from the environment and fixes their location within a tissue, plant cells are responsive to mechanical stimuli. However, several questions remain unanswered around how mechanical stimuli are perceived and translated into cellular responses.

Here we establish a system enabling application of quantifiable localized mechanical stress while simultaneously capturing cellular responses with high spatio-temporal resolution using confocal imaging. We show that this system enables estimation of locally applied pressure and provides access to the temporal and spatial details of subcellular events in intact living tissues upon touch. We propose that observing these subcellular events at high spatiotemporal resolution and linking their dynamics to the intensity of mechanical stimuli may uncover the molecular mechanisms underlying plant cell responses to touch.

## Introduction

Plant cells are able to respond to a variety of mechanical stimuli, and developmental patterns of mechanical stress are interlinked with cellular processes. For example, in development, cell-division plane orientation (Kny, 1896; Lintilhac and Vesecky, 1984; Louveaux et al., 2016a) and localization of proteins have been linked to tissue mechanical stress patterns (Bringmann and Bergmann, 2017; Elliott et al., 2024; Heisler et al., 2010; Nakayama et al., 2012; Piepers et al., 2025). Similarly, response to intrinsic mechanical stresses arising during tissue growth has been tightly linked to developmental processes at the organ scale, such as organ shape maintenance (Hervieux et al., 2017), lateral root emergence (Vermeer et al., 2014) and directional growth (Nemec-Venza et al., 2026). With regards to responses to external mechanical stimuli, the striking fast movement of whole organs in some plants upon touch has been known for centuries (Darwin, 1875; Ellis, 1770; Hooke et al., 1665) and, more recently, details of signalling events linked to these movements have been described (Hagihara et al., 2022; Procko et al., 2022; Suda et al., 2025, 2020). Calcium signals are one of the most studied fast cellular responses upon touch (Bellandi et al., 2022; Howell et al., 2023; Matsumura et al., 2022; Pantazopoulou et al., 2023), but are not the only ones. Release of elicitors (Bellandi et al., 2022), endoplasmic reticulum clustering (Hardham et al., 2008), actin clustering (Branco et al., 2017; Hardham et al., 2008) and microtubule reorganization (Hardham et al., 2008; Jacques et al., 2013) all have been shown to be triggered by touch.

Several cellular responses and downstream signalling events occurring after external stimuli like touch have been studied (Bellandi et al., 2022; Branco et al., 2017; Hardham et al., 2008; Howell et al., 2023; Jacques et al., 2013), however in most cases the molecular mechanisms that allow perception of mechanical stress and subsequent orchestration of the observed cellular responses still remain to be elucidated. For example, the mechanisms behind the endoplasmic reticulum and actin behaviour upon touch, and the mechanism of elicitor release upon touch still remain unknown. We propose that the molecular pathways linking touch perception to these fast cellular responses could be elucidated by overcoming a main limitation common to most standard methods: the difficulty in monitoring the magnitude of the local stress while simultaneously performing live correlative microscopy at sub-cellular level. These responses develop within seconds to minutes of stimulus, and their dynamics and spatial patterning around the touch site reflects in part the molecular machinery they rely on. Thus, live confocal imaging at sub-cellular resolution combined with a fine control of simultaneous local mechanical stimulation will help shed light on the molecular mechanisms behind these cellular responses.

Plant anatomy poses significant challenges for confocal-level live imaging of cellular responses upon touch in intact tissues, due to often non-transparent organs composed of walled cells usually organized in multilayered tissues. Ablation of cells (via laser or sharp needles) and osmotic stress have been used extensively to deduct effect of mechanical stress on cellular and tissue processes (Bringmann and Bergmann, 2017; Hamant et al., 2008; Hoermayer et al., 2020; Nakayama et al., 2012; Sampathkumar et al., 2014; Serra et al., 2026), however these methods introduce other variables alongside mechanical stress – for example extensive release of damage-associated molecular patterns due to ablation of cells, and alteration of osmotic potential and ion levels of the extracellular environment. Mechanical stress application at the whole tissue level via folding (Serra et al., 2026) or quantifiable stretching (Robinson et al., 2017; Trozzi et al., 2025) has shed light on mechanical properties of tissues and dependency of developmental processes on stress patterns. However, with these methods the stimulus is applied across the whole tissue rather than locally and the coupling with live confocal imaging poses specific challenges for these systems. Similarly, tissue-wide compression has highlighted interesting patterns of microtublule response (Louveaux et al., 2016b; Sampathkumar et al., 2014) and ER-contact site behaviour (Perez-Sancho et al., 2015), but did not allow for a fine time-resolution of the dynamics of such processes nor for studying responses to localized stress application. The use of nano-indenters (Beauzamy et al., 2015; Branco et al., 2017; Louveaux et al., 2016b) and cellular force microscopy (Majda et al., 2019) has allowed for quantifiable local mechanical stimulation, but did not allow for simultaneous confocal imaging. Similar limitations are shared by AFM-based systems – while they open to remarkable opportunities for quantification of tissues mechanical properties and applied forces, their application remains challenging when it comes to the need to couple them with simultaneous confocal imaging at sub-cellular resolution in thick and multilayered tissues. Overall, several tools for mechanically stimulating and imaging plant tissues are available, but a minority allow for localized application of mechanical stress and simultaneous imaging. Recent work demonstrates the power of quantifiable indentation-based local mechanical stimulation coupled to confocal microscopy to elucidate rapid dynamic responses to stress in the form of intracellular calcium signals (Howell et al., 2023). However, the application of these kind of tools remained so far limited to tissue-wide observations and quantification of applied forces rather than pressures.

Here we present a system that allows localized stimulation, estimates of the locally-applied pressure, and simultaneous live confocal imagining at the touch site with sub-cellular resolution in intact, living, multilayered tissues. We propose that using this set-up to systematically study details and dynamics of cellular responses can shed light on the molecular mechanisms underlying plant cells perception and response to touch.

## Methods

### Needle manufacturing

Needles were pulled from Drummond microcaps 50 µl glass capillaries using a Narishige PC-10 pipette with settings as described in the text. The sharp tip of the needles was then melted using a microforge (Alcatel) to create hemispherical blunt needle tips. For quality control, needles were then imaged using a Nikon SMZ18 stereomicroscope equipped with an Orca flash 4 Hamamatsu C11440 camera. The diameter of the tip was measured in Fiji by manually fitting a circle to the tip of the needle.

### Needle calibration and estimation of pressure

The load cell used for needle calibration was Futek LSB200 (20g, model FSH03868) and was set up on the blueprint of the ACME system (Robinson et al., 2017), with a LabView based software (software version used 4.1.1, (Smithers et al., 2026)). The software allowed to access the readout of the applied force from Futek load cells at 100ms time interval in real time. Readout of the load cell relies on the Futek-provided DLLs as well as on its factory calibration. The needle was placed at the closest distance possible from the mobile plate of the load cell where the load cell did not register changes in force, then, by performing defined displacements of micromanipulator motor, the needle was moved vertically to push on the mobile plate of the load cell. The needle was then lifted and lowered again after a few seconds interval to create repeated touches on the load cell mobile plate. Force readouts from the software were then analyzed using an R (R Core Team, 2025) script (publicly available at the Github repository https://github.com/AnnalisaBe/Microindentation-toolbox.git). For pressure estimates, the plant-needle interface was approximated to the curved surface of a half sphere. The pressure applied on the tissue upon needle movement was then estimated as the force exerted by the needle movement (as calibrated by the load cell) divided by the curved surface of a half sphere with the same radius as the needle tip. The angle of the needle holder was kept constant between calibration steps and experiments.

### Micromanipulator and micromanipulator software

Micromanipulator motors and controller with associated software were purchased from Sutter (MPC-325 system, with MPC200 controller and two PC225 micromanipulators).

### Plant material

Seeds of Arabidopsis thaliana ecotype Col-0 expressing RFP-HDEL as endoplasmic reticulum (ER) makerk originated from Brault *et al*. (2019).

### Plant growth

Seeds were sterilized for 5 min in 70% ethanol, then rinsed with sterile water three times. After 3 days of stratification in the dark at 4°C, seeds were sown on 0.5X Murashige Skoog media with vitamins, without sugar. Plates for hypocotyls experiments were placed for 2 hours in the light at room temperature, then in the dark in vertical position at 22°C for 3 days.

### Plant samples handling and imaging

For mounting the samples, a thin layer of Kwik-Sil was deposited on a glass slide using a glass coverslip as a spatula - inclining the glass coverslip at 45 degrees angle and dragging it along the length of the slide allowed for the distribution of an even layer of glue. The glue was let cure for 7 minutes, then a sample was carefully lifted from the plate and placed on the glue, and water was added immediately. Samples were let recover for at least an hour before imaging.

Imaging was carried on a Zeiss LSM980, using a water-dipping objective Zeiss N-Achroplan 40x/0.75 with working distance of 2.1 mm. Excitation wavelength was 561 nm, RFP emission was collected with a Spectral detector between 570 nm and 640 nm. Transmitted light images for monitoring needle movement and position were collected with a T-PMT detector.

### Image analysis and quantification

Images were quantified in Fiji. In order to collect the whole surface of the curved tissue, the files are collected as Z stacks. To estimate timings, times of each optical slice of the Z stacks were sourced manually from the metadata. To measure fluorescence increase after touch, a Z projection obtained from the sum of the optical slices in the Z stack was used. On this projection, the average intensity in the area under the needle after the needle was lifted and in a control untouched area of the same cell were collected by two hand-drawn ROIs of the same size and shape. The ROI placed in the untouched area of the cell was used as reference ROI. The average intensity in the reference ROI was subtracted from the average intensity in the touched ROI, the result was then divided by the average intensity in the untouched ROI. This ratio was used as a proxy for ER accumulation at the touch site. For this analysis a single time point was chosen, corresponding to the time point after the needle was lifted when the signal was the most intense. For analysis of the signal accumulation over time, a line of thickness 50-100 pixels was drawn along the length of the touched cell and centered at the touch site (as shown in Fig. 5A). A semi-automated Fiji (Schindelin et al., 2012) macro and R (R Core Team, 2025) script were used to analyze these data and calculate average profiles of the signal at each time point. Codes are available on the Github repository https://github.com/AnnalisaBe/Microindentation-toolbox.git. Briefly, the fluorescence intensity of pixels at the same position along the line were averaged to create a separate profile of the signal at each time point and maintain the positional information of the intensity along the line. Ggplot2 (Wickham, 2009) geom_smooth function was used to fit a loess curve on the experimental datapoints. For quantification of the ER network, a single optical slice at the cell surface was considered and the image was analyzed in Fiji using pre-existing functions. Briefly: the area of interest showing clear ER network was cropped from the original image, the multipoint tool was used to mark the center of each area delimited by RFP-HDEL network strands, the points were added to the ROI manager, the points were automatically drawn as white dots on a new image with the same size of the analized area but with black background, the Voronoi function in Fiji was used to split the binary image into regions using lines equidistant to the borders of the two nearest dots, a mask was created using the lines generated by the Voronoi function, the Analyze Particles function of Fiji was used to measure the area of the Voronoi regions. The area of the Voronoi regions in the touch zone was compared to the area of the Voronoi regions in the zone not interested by touch, before touch and after touch. Statistical comparisons and plots were done in R (R Core Team, 2025)

## Results

### Assembly of the technical platform: micromanipulator coupled with confocal microscope

As our interest resided in observing sub-cellular details of the response to touch around the touch site, we aimed to image and touch the sample from the same side, to exploit the full imaging potential of the equipment at hand. In order to be able to mechanically stimulate the sample and image directly on the site of stimulation, we built the system around an upright confocal microscope equipped with a long-working distance objective and coupled with a commercial micromanipulator set-up from Sutter. A support tower holding a cantilever platform and bolted on the microscope antivibration table enabled us to adjust the height of the micromanipulator above the height of the microscope stage (Fig 1A), allowing the microscope stage to move independently from the micromanipulator not to be subjected to the weight of the micromanipulator motors. To reach the sample, glass needles manufactured in-house were mounted on the micromanipulator arm via a long commercial needle holder. The sample mounting required stabilizing the sample without minimizing the mechanical stress derived from the mounting set-up. Upon testing different sample mounting methods, including partial embedding in low melting point agarose and different adhesives, we found that the non-fluorescent veterinary glue Kwik-Sil provided the best compromise for sample immobilization, minimization of autofluorescence and sample viability. In order to keep the sample from drying and allowing imaging with a water-dipping objective, a small chamber was created around the sample using vacuum grease (Fig 1B). Unsing this system, the sample could be touched and imaged on the same side before, during and after the mechanical stimulation.

**Fig. 1:**
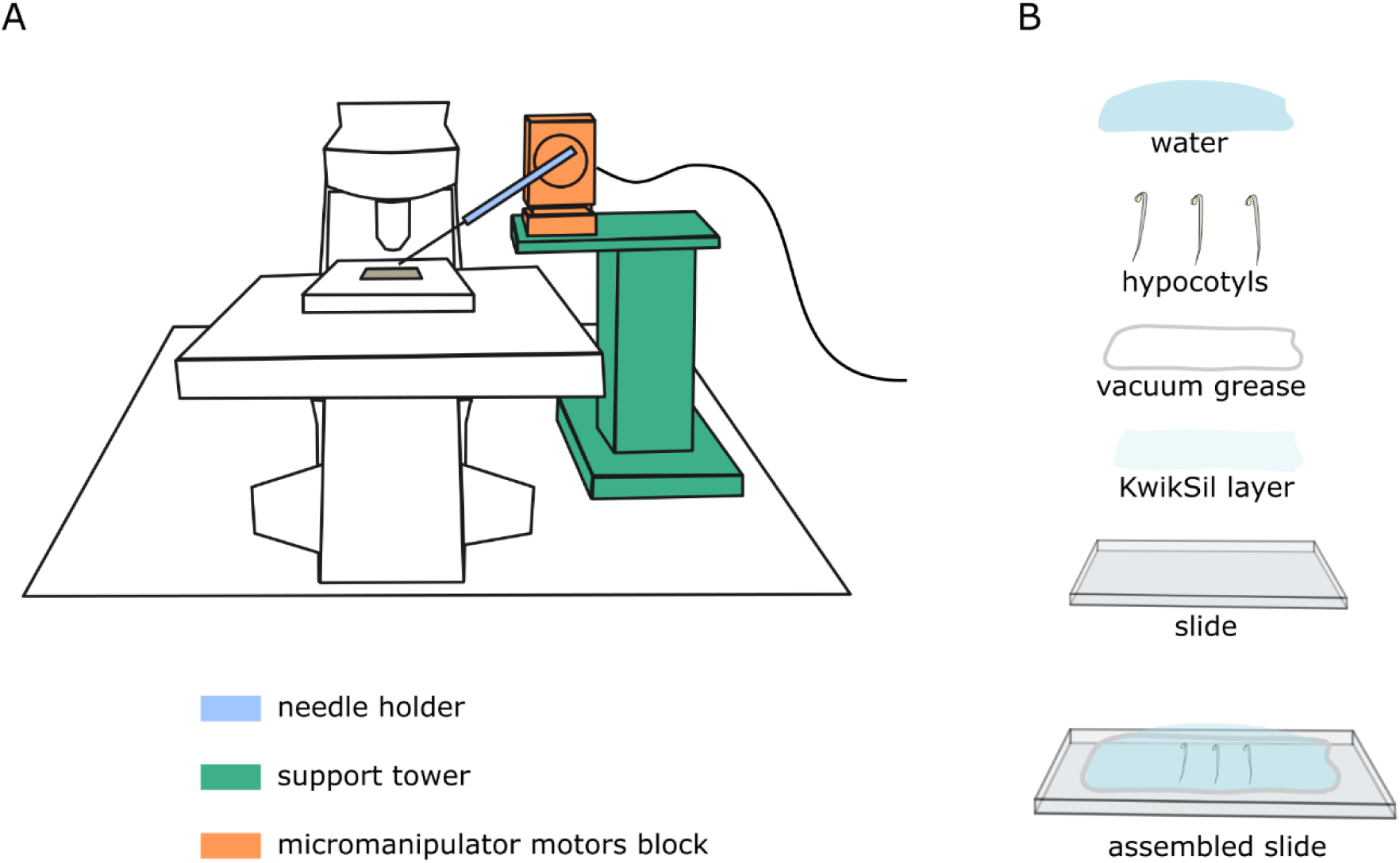
Schematic of the experimental set-up. A) Schematic of the Sutter micromanipulator coupled to an upright confocal microscope. The piezo motors of the micromanipulator block (orange) are held by a support tower and a cantilever platform (green) above the microscope stage. A long needle holder arm (cyan) allows reaching the sample. B) Schematic illustrating the details of the sample mounting. A think layer of Kwiksil glue holds the sample in place, a perimeter of vacuum grease holds water to form a pool.

### Manufacture of calibrated needles with hemispherical tips

As a tool for applying mechanical stimuli to the tissue we opted for in-house manufactured glass needles. Although a variety of needles of different materials and sizes are commercially available, we reasoned that the glass needles would allow us maximum flexibility and adaptability in terms of tuning the size and shape of the tool. While glass needles can be easily manufactured from glass capillaries by the use of a pipette puller, the resulting tip is sharp and easily leads cell rupture. To minimize the chance of accidental cell rupture upon touch, we used a microforge to create hemispherical needle tips of controlled sub-cellular size. Using this device, under an optical binocular microscope, a sharp glass needle is guided via the use of micrometric knobs towards a current-heated platinum wire. The shape and size of the needle tip can then be controlled by adjusting angle and distance of the needle from the wire, as well as wire temperature and time of exposure of the glass to the heat. Varying needle pulling parameters and microforge parameters we were able to obtain needles with rounded tips and different sizes (Fig. 2). Higher temperatures (and/or lighter weights) create long slowly tapering tips where the lateral walls of the tip proceed forward in near parallel fashion, lower temperatures (and/or heavier weights) create a tip with a faster tapering degree that ends in a sharper tip. Needles created with needle puller temperature setting 60 and 4 weights were chosen for the rest of this work, as they allowed to easily achieve rounded tips of diameters spanning from 8 µm to 20 µm, optimal for studying the response to touch and minimizing accidental cell ablation, while maintaining a relatively small sub-cellular size footprint.

**Fig. 2:**
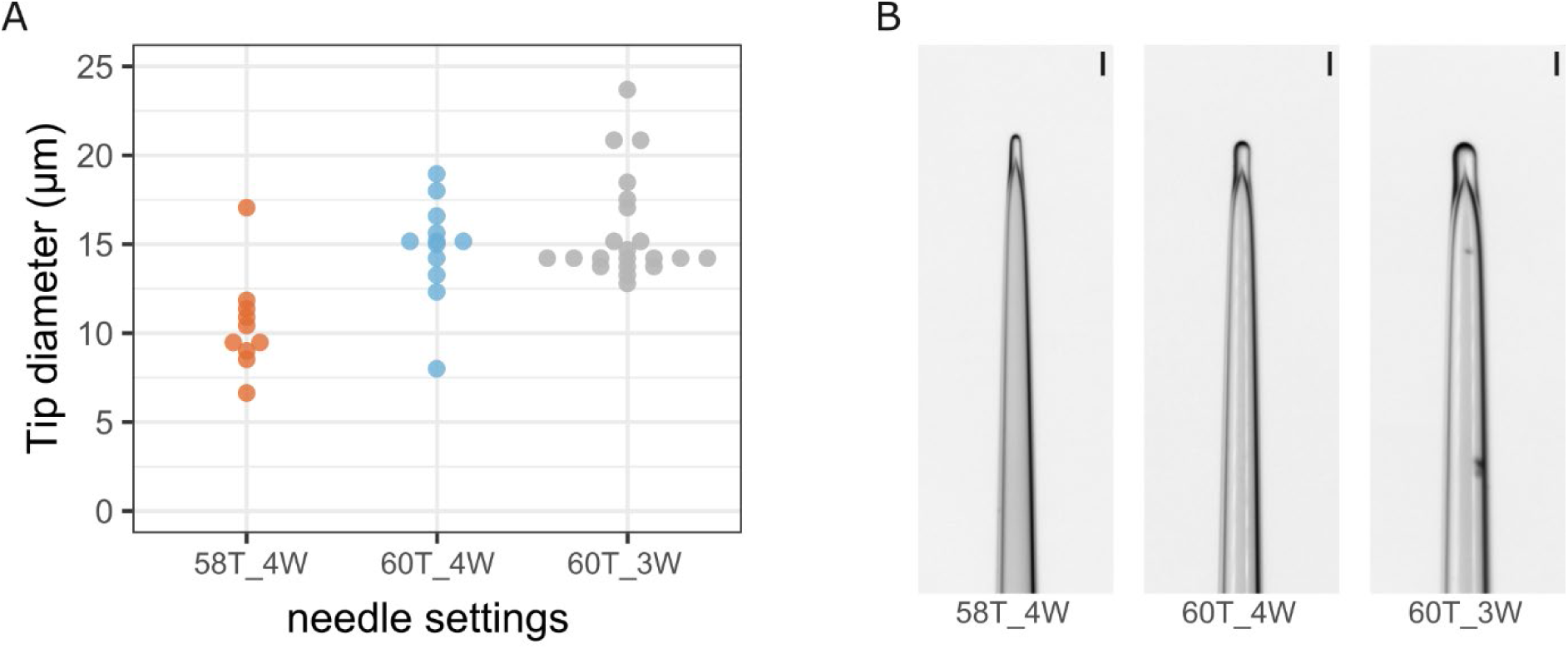
Size and shape of the in-house manufactured blunt glass needles. A) Tip diameter for blunt needles created using three different needle puller settings. Adjusted settings for the needle puller comprise a temperature step control (T) to tune glass melting, and a number of weights (W) to tune the pulling force. B) Microscopy images of glass needles pulled with the three different needle puller settings and blunted using the microforge, achieving different tips diameters. Scale bars 20 µm.

### Calibration of the indentor

Glass micropipettes of calibrated flexibility have been used before to estimate forces (Backholm and Bäumchen, 2019; Howell et al., 2023; Popowski et al., 2025). In order to estimate the level of mechanical stress applied on to the sample, we calibrated the needles exploiting a load cell placed in vertical position (Fig. 3A). The load cell was secured on a support tower and bolted on a fixed plate, while another plate was secured on the top mobile part of the load cell. Via the use of the provided Sutter software, the micromanipulator was programmed to repeatedly move the needles of 5, 10 and then 20 µm towards the load cell starting from the point of first contact with the top mobile plate. As a result, the load cell read-out appeared as repeated peaks of force surge over the baseline (Fig. 3B). Each needle was tested over 3 repeated test cycles, where one test cycle is composed by 10-12 pushes of 5 µm, followed by 10- 12 push of 10 µm, followed by 10-12 push of 20 µm. Analysis of the load cell read-out across all the cycles and needles movement allowed to quantify the magnitude of the stimulus associated to a defined needle movement for each needle (Fig. 3C).

**Fig. 3:**
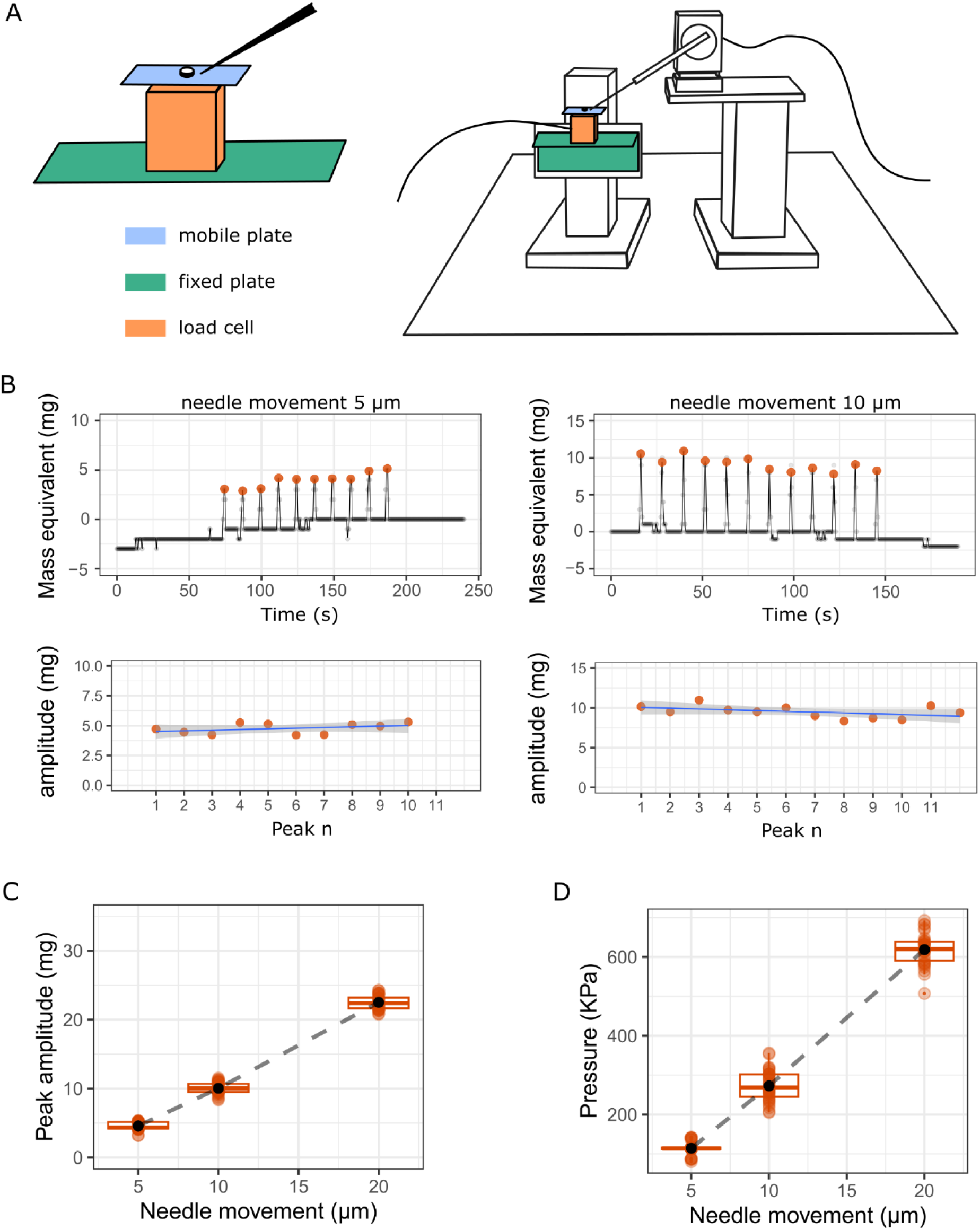
Needles calibration. A) Simplified schematic of the load cell and micromanipulator set up used for the calibration of needles (right) and detail of the load cell – needle contact (left). The load cell (orange) is mounted in vertical position on a fixed plate (green) bolted to a support tower. The needle guided by the software pushes on a mobile plate (blue) secured on the mobile part of the load cell. B) Load cell output (top) and quantification of the peaks amplitudes (bottom). Each needle movement is read in the load cell output trace as a peak. In the load cell output traces, grey dots are load cell output datapoints, dark grey line is loess curve fitted on the experimental data, orange dots are detected peaks. In the plots for peak amplitude quantification (bottom), orange dots are peak amplitude relative to the baseline, blue line is a straight line of best fit generated using a linear model, shaded area shows the confidence interval for the line. C) Detected peaks amplitudes in response to needle movements of 5, 10 and 20 µm in 3 subsequent testing cycles. Orange dots are experimental data points, black dots indicate the mean value of the group, dashed line connect means across groups. D) Pressure applied on hypothetical tissue, estimated considering the experimental data in C, the needle diameter (15 µm) and a needle-tissue interface approximated to a half sphere of diameter 15 µm.

While it is temping to compare the force exerted by the needles on the load cell to the weight of insects for example, it should be taken into consideration that the exact pressure applied on the sample depends not only on the force but also on the needle-sample interface. The complex topology of plant samples does not allow for an exact calculation of the sample-needle area interface and the precise incidence angle between the needle and the tissue. Thus, we propose a proxy for pressure applied on the samples, based on the needle size and the load cell force output (Fig 3D), that is specific for each needle upon defined movements (in Z axis) keeping the needle angle constant. For the needle taken as example, considering its diameter (15 µm) and the force measurements of the load cell, we estimate that a movement of 10 µm would apply locally ∼270 KPa on the tissue (Fig. 3D).

### ER quickly clusters at touch site

Previous studies reported that ER quickly clusters at the touch site (Branco et al., 2017; Hardham et al., 2008), but live confocal imaging and estimation of the stimulus intensity were only achieved separately, not at the same time. We decided to test if we could observe ER accumulation at the touch site with our system, and if we could reveal more details of the ER features and accumulation dynamics by performing simultaneous live imaging and quantifiable stimulation. Hypocotyls were chosen for this experiment as it is a tissue in active growth naturally exposed to mechanical cues and able to quickly respond to stimuli (Branco et al., 2017; Lindeboom et al., 2013). We tested ER accumulation with needles of either 8 µm or 15 µm diameter, combined with different ranges of movement and length of stimulus (Fig. 4). Modulating the range of movement in the Z axis with needles of different diameters, we can modulate the pressure applied on the sample. With movements of 10, 15, and 20 µm we covered a range of estimated pressures from ∼200 to ∼2000 KPa (Fig. 4A). We quantified fluorescence intensity level increase in the touched area after touch, compared to a control untouched area in RFP-HDEL etiolated hypocotyls. In elongated cells located 500-1000 µm from the apical hook of etiolated hypocotyls, we found that ER clusters at the touch site across the whole range of pressures tested (Fig 4A). We also asked if the ER accumulation was dependent on a maintained stimulus, and to test this we applied pressures for different durations. We tested stimuli durations between ∼2 s and ∼700 s, and we observed that accumulation of ER at the touch site was triggered in all the tested stimuli (Fig. 4B), indicating that prolonged touch is not necessary to trigger the response.

**Fig. 4:**
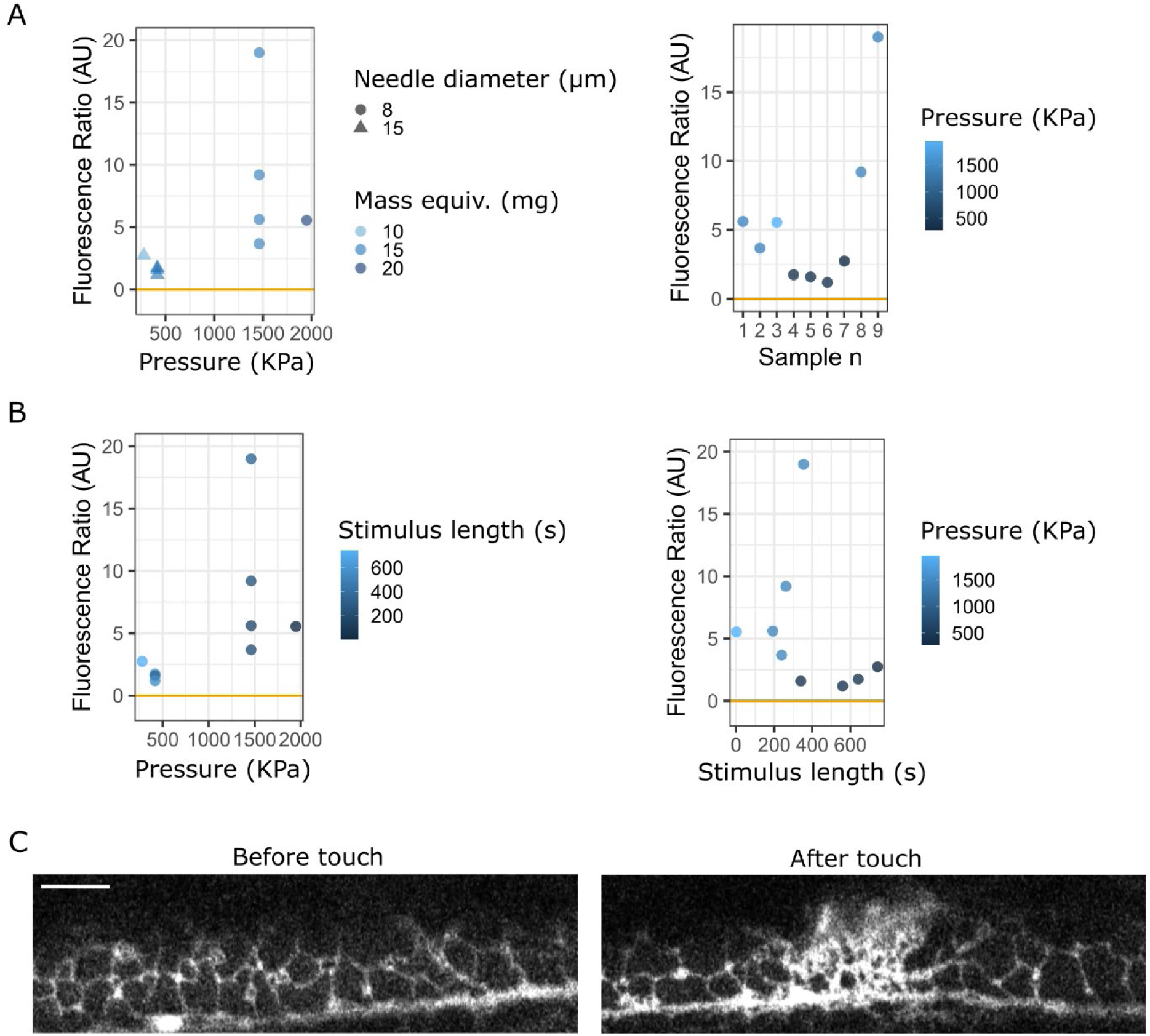
ER accumulation upon touch. A) Fluorescence increase at the touch site after touch, in relation to the pressure applied. B) Fluorescence increase at the touch site after touch in relation to the length of the applied touch stimulus. In A and B, Fluorescence ratio is average intensity increase at the touch site divided by average intensity in untouched control area. Horizontal orange line corresponds to value of 0, indicating no increase in fluorescence before and after touch. A and B are based on the same datasets, different plots shown to examine the response in relation to different variables. C) Confocal image showing the surface of a single hypocotyl cell in a single imaging plane before touch (left) and right after the needle was lifted (right) in etiolated hypocotyl epidermal cell. Scale bar 10 µm. Fluorescence signal is RFP-HDEL.

Quantifying the ER accumulation in the Z projection allows us to capture the accumulation in the whole volume and along the curvature of the cell, but the details of the ER network at the surface of the cell go lost in the projection. To visualize structural details of the ER network, we analysed a single optical section at the surface of the cell. While it is not possible to reliably image the cell surface under the needle during touch, we were able to observe structural changes in the ER network just after the needle was lifted compared to the structure observed in the same area before touch. After a prolonged touch of ∼ 600 s with a needle of 15 µm diameter applying a pressure of ∼400 KPa, the ER in the area that was under touch site appeared as a denser network immediately after lifting the needle (Fig. 4C, Fig 1S). This also reveals that the ER accumulation area (covering a width of ∼20 µm) is slightly larger than the needle (15 µm diameter).

Next, we investigated the dynamics of ER accumulation over time. For this, we analysed the average profile of RFP-HDEL signal along the length of the cell over time (Fig 5). Observing the RFP-HDEL signal accumulation around the needle edge during touch confirmed the observation that the ER accumulation area is larger than the needle contact area (Fig 5A, Fig. S2). By comparing the profile of the signal during touch and over time in the touched area after the needle was lifted, we noticed that the signal seemed to present two phases of accumulation – one during touch, and a renewed accumulation starting when the needle was lifted (Fig. 5A-B, Fig S2) - the accumulation of ER during touch is visible in the rising profile at the edge of the needle (Fig. 5A, Fig. S2), the peak visible immediately after the needle was lifted (Fig. 5B, middle panel) then develops in a peak of higher intensity in the 80 s following the lifting of the needle (Fig 5B right panel). To overcome the limitation of the needle obstructing the view and visualize the dynamics of ER accumulation in the touched area with a higher temporal resolution, we studied the ER accumulation upon a fast stimulus (2 s touch) and monitored the intensity of the RFP-HDEL marker along the length of the cell (Fig. 5C). This revealed that a short stimulus is sufficient to trigger ER accumulation, and that the ER accumulation continues when the needle is lifted - the intensity profile, centered on the needle central touching point, develops in a more intense peak as time passes (Fig. 5C). The area of ER accumulation upon quick touch and release also expands beyond the area directly touched by the needle, like the ER accumulation upon prolonged touch (Fig 5B-C). Overall, this suggests that, rather than the maintained pressure, the signalling events triggered by touch/release could be at the base of ER accumulation.

**Fig. 5:**
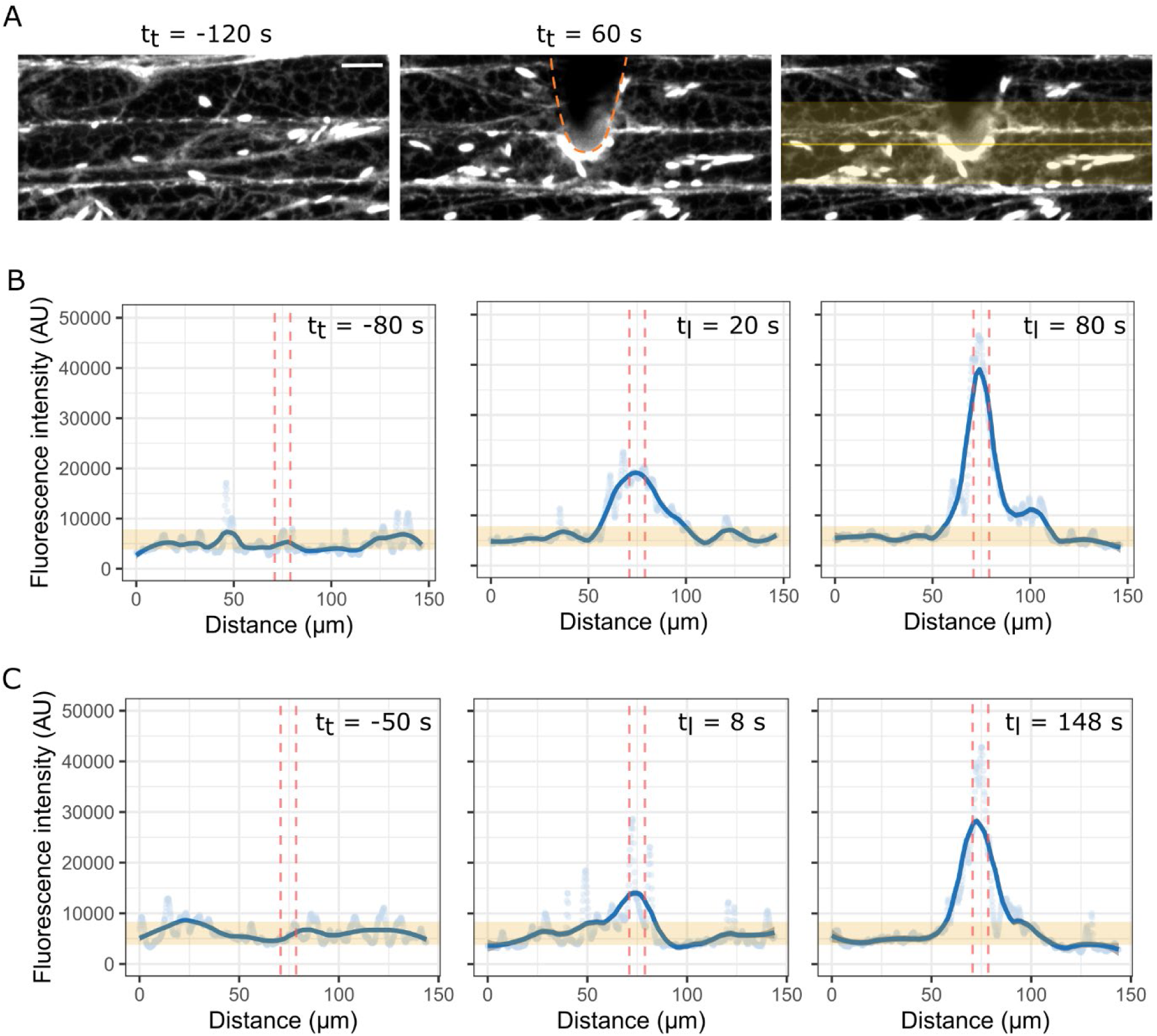
Dynamics of ER accumulation upon touch. A) Confocal microscopy images showing projections (sum of slices) of hypocotyl epidermal cells before and during touch. Needle outline is indicated by orange dashed line. In the right image, yellow shaded area and yellow line indicate the region of interest used to create the profile plots over time. Scale bar 10 µm B and C) Profile plots of the average intensity over time in two different samples. With reference to the region of interest marked in panel A right image, the distance in the profile plots corresponds to µm along the yellow line, and the profiles are created by averaging pixels in the yellow shaded area that are positioned at the same longitudinal distance along the line. Orange shaded ribbon represent baseline fluorescence +/- SD. Blue dots represent experimental data of measured average profile. Blue line represents loess fitted curve to describe the profile. Vertical red dashed lines demark the area touched by the needle. B are profiles over time of ER response to prolonged touch of ∼300 s, with a needle of 8 µm diameter, exerting a pressure of ∼ 1400 KPa. C are profiles over time of ER response to a quick touch of ∼2 s, with a needle of 8 µm diameter, exerting a pressure of ∼ 1900 KPa. Across the figure, t_t_ corresponds to time in relation to the start of the touch event, t_l_ indicates time from the moment the needle was lifted.

## Discussion and Conclusions

Here we developed a system that allows localized quantifiable mechanical stimulation and simultaneous confocal imaging at the stimulus site, with high spatial-temporal resolution. Manufacturing and calibrating the needles allows to create needles of different size and stiffness, in turn opening the road to study the effect of different variables on cellular responses of interest. It could for example be of interest to systematically test the effects of increased pressures and different curvatures of the membrane on cellular responses. Moving forward with this system, however, it is important to keep in mind that the pressure quantification method proposed here allows only an estimation of the pressure and not an exact calculation, since the exact pressure applied locally on the cell may be affected by other factors such as the topology of the tissue which dictates the incidence angle and contact interface at a microscopic level. Finally, while we are focussing on the applied pressure, the perceived signal for the plant may be a different one (for example displacement or alteration of the membrane-wall continuum), and considerations on the perceived signal and its magnitude in relation to the applied pressure may need to take into account the fact that the tissue is pressurized (Beauzamy et al., 2015; Vella et al., 2012). Overall, while the exact pressures applied on the tissues cannot be calculated with the simple formulas used here, we propose a method for approximate applied pressure estimation starting from calibrated needles of defined size.

We showed ER response to touch upon application of 5-20 mg weights which, based on our calculations, would result in estimated pressures in the range of ∼200 KPa to ∼2000 KPa at the needles tips. For reference, car tires are usually inflated to 220-250 KPa pressure levels. A 10 mg weight is comparable to the weight of a common housefly *Musca domestica* (10-15 mg) (Goulson et al., 1999). In the case of a fly this weight is divided over six feet, each having two pads each of diameter ∼100 µm (Niederegger and Gorb, 2003). Dividing the ∼12 mg weight over 12 pads of ∼100 µm each, each pads applies a pressure of ∼0.6 KPa. While each pad also presents a sharp claw that could itself apply more localized (and possibly higher) pressure on the tissue during movement (Niederegger and Gorb, 2003), the division of the weight over several contact points could still result in lower pressures than the ones applied here with the glass needles. While the lower pressure we tested was ∼200 KPa, we expect that there could be cellular responses at even lower pressure. As the textbook image of the cell wall is the one of a stiff load-bearing barrier, it would be of interest in the future to test the minimal pressure that can cause cellular responses, for example investigating if and what cellular responses can be observed upon pressures in the range of ∼1-0.5 KPa. However, plant tissues do experience in nature comparable pressures to the ones tested here – for example, the necrotropic fungus *Botrytis cinerea* can exert a penetrating pressure of 500 KPa (Müller et al., 2024) and the turgor pressure of the appressorium of the rice blast fungus *Magnaporthe oryzae* has been estimated at 8000 KPa (Howard et al., 1991). Given the dependency of the estimated pressure on needle size, movement and flexibility, this system can therefore be tuned and exploited to test the range of pressures experienced by plant tissues in nature, from contact with insects to attach of pathogens.

Localized stimulation and simultaneous confocal imaging at sub-cellular resolution with direct visualization of the stimulus site makes this system suitable to image non-transparent and thick samples. Further, this set-up opens the road to study responses upon touch with high spatio-temporal resolution, allowing the dissection of their dynamics and their structural features. Consistent with the work of Hardham and colleagues (Branco et al., 2017; Hardham et al., 2008), here we reported the fast accumulation of the ER at the touch site. While we could observe ER accumulation within 8 s of touch, it is possible that the ER accumulation starts even faster but our imaging set-up and settings did not allow us to detect that. Time-lapse imaging before, during, and after touch allowed us to quantify the dynamics of ER accumulation upon prolonged or short touch. Interestingly, we observed that, during prolonged touches, there seem to be a first burst of accumulation during the phase of pressure application followed by a second burst of accumulation after pressure release. This is reminiscent of the two-phased calcium signal reporter by Howell et al. (Howell et al., 2023) upon application and release of pressure, and could suggest a link between secondary signals triggered upon touch and the touch response observed here. We also report that the ER appears as a denser network at the touch site upon a prolonged touch. Further study of the spatio-temporal dynamics of the responses upon touch and structural development of the structures of interest over time has the potential to shed light on molecular mechanisms at the base of this and other touch-triggered cellular process.

## Data accessibility

Additional data and information can be provided upon request. Scripts used in this work are available at https://github.com/AnnalisaBe/Microindentation-toolbox.git.

## Author contributions

A.B. conceived and designed the project, developed methodology, developed technical platform, acquired the data, analysed and interpreted the data, wrote the initial draft, acquired funding. C.L. provided technical assistance to acquire and assemble the technical platforms. C.L., N.G. and C.K. set up the load cell system. M.O.L. developed and provided the load cell readout software. D.A. acquired data. A.B., C.K., I.F.L. and O.H. acquired funding. All authors provided feedback on the initial draft.

## Acknowledgements

We thank Simone Bovio (ENS Lyon) and Juan Alonso Serra (University of Helsinki) for discussions and advice during the development of the technical platforms.

Funding sources: EMBO Long Term Postdoctoral Fellowship ALTF_81-2023 (A.B.), ANR ROCnROS and ANR-21-CE13-0049 (I.F.L.), ERC-2020-Stg 948514—EDGE-CAM (C.K.), ERC-2021-AdG-101019515 “Musix” (O.H.), M.O.L. is supported by the Gatsby Charitable Foundation as part of the Sainsbury Laboratory Cambridge University microscopy core facility.

