## Supplementary_information for "Live confocal imaging of cellular touch responses upon local quantifiable mechanical stimulation in plant cells" (from another version of this preprint: version 1, 2026-09-13)

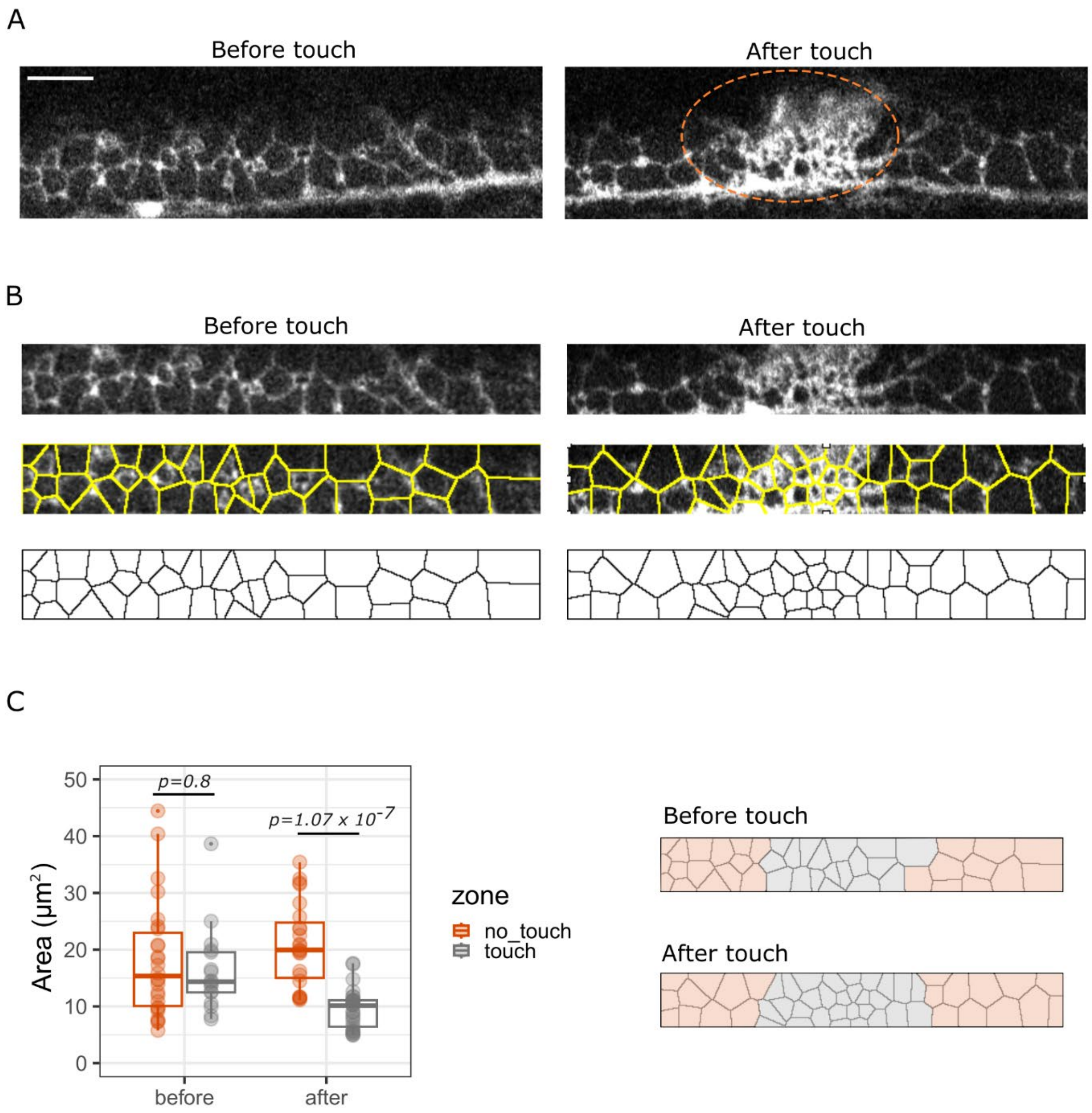


**Fig S1: Quantification of ER network upon touch.** A) Confocal images of a single optical section of the cell surface of hypocotyl cell expressing RFP-HDEL, as in Fig 4C. The area interested by the touch-triggered response is marked by the orange dashed line. B) Steps in the quantification pipeline explained in the methods section, for the same area before touch (left) and after touch (right). The area of interest was cropped from the image in A (top), mask of Voronoi regions identified as explained in the methods overlayed over the microscopy image (middle), mask of the Voronoi regions alone (bottom). C) Plots showing the area of the Voronoi regions before and after touch, in the zone interested by the touch and in the zone not subject to touch. On the right, colour-coded masks show the touch/no-touch zones (indicated by colours following the plot legend). The ‘touch’ zone before touch indicates the approximate area that will be subject to the touch response after touch. Statistical comparison was conducted with Wilcoxon rank-sum test, resulting p values are indicated in the plot.

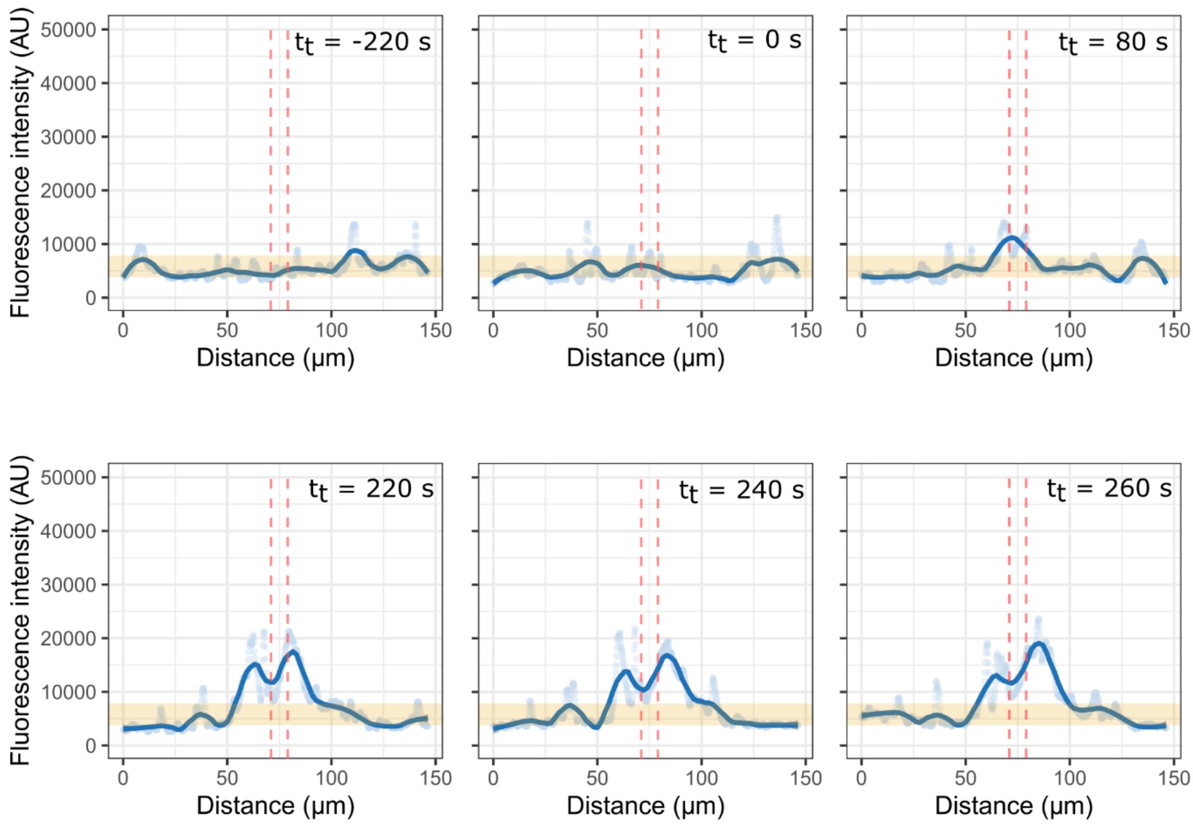


**Fig S2: Details of ER accumulation during touch.** Plots with same details as in Fig 5B, based on the data in Fig 5A, showing detail of the ER accumulation during touch, before the needle is lifted. Orange shaded ribbon represent baseline fluorescence +/- SD. Blue dots represent experimental data of measured average profile. Blue line represents loess fitted curve to describe the profile. Vertical red dashed lines demark the area touched by the needle. The experiment involved a prolonged touch of ~300 s, with a needle of 8 μm diameter, exerting a pressure of ~ 1400 KPa. Across the figure, $t_t$ corresponds to time in relation to the start of the touch event.
